# Sex differences in the developing human cortex intersect with genetic risk of neurodevelopmental disorders

**DOI:** 10.1101/2025.09.04.674293

**Authors:** Kelsey Hennick, Yang Sui, Druha Karunakaran, Ally Nicollela, Rachel Leonard, Rebecca Meyer-Schuman, Hanna Berk-Rauch, Tianyun Wang, Aravinda Chakravarti, Huda Y. Zoghbi, Evan E. Eichler, Tomasz J. Nowakowski

**Author notes:** These authors contributed equally (Kelsey Hennick, Yang Sui).

## Abstract

Autism is highly heritable and diagnosed more frequently in males than females. To identify neurodevelopmental processes that might present sex-biased vulnerability, we generated transcriptomic and epigenomic profiles of cell types present in the prenatally developing human cerebral cortex of 27 males and 21 females. By intersecting sex-biased molecular signatures and genes with *de novo* mutations in male and female autistic probands, we reveal two points of vulnerability contributing to the sex-biased penetrance in neurodevelopmental disorders (NDDs). First, we show that NDD risk genes are biased towards higher expression in females, identifying the NDD gene *MEF2C* as a critical transcription factor for female-biased expression. Second, we identify a significant contribution of X chromosome genes to NDD pathobiology. We construct a gene regulatory map of X-linked risk genes to enable functional studies of genetic variants that likely disrupt gene expression in the developing brains of autistic males. Together, these results point towards an outsized contribution of the X-chromosome to both the origin of sex differences in the developing human cortex and NDD vulnerability. We propose a model where female-biased vulnerability is driven by coding variation within genes while male-biased vulnerability is driven by noncoding variation in regulatory elements that affect gene expression.

## Introduction

Development of the human cerebral cortex is highly dynamic, and control of cortical gene expression across developmental periods, cell types, and individuals requires extensive networks of transcriptional regulators, and thousands of ‘regulatory’ genomic loci. The prenatally developing midgestation cerebral cortex is highly enriched for co-expression of high-confidence autism risk genes identified through studies of *de novo* protein-damaging variants^1–7^, particularly within deep cortical layer neurons^1,5,6^. While studies of gene expression and epigenomics have begun to uncover the blueprint of gene regulatory networks that give rise to relevant brain regions and key cell types^8–12^, how these processes vary between males (XY individuals) and females (XX individuals)^13–15^ has not been extensively examined. This represents an important gap relevant to our understanding of the etiology of autism, which shows a significant bias towards male diagnosis^4,16,17^.

Bulk tissue measurements of gene expression in male and female brains have emphasized the contribution of chromosome X- and Y-linked genes^18^. Consistent with this observation, X-chromosome-linked *de novo* gene mutations are overrepresented in females, likely due to the buffering effect of two copies of the X chromosome and hypothesized lethality in males^3^. Single-cell studies have revealed autosomal differences, with most differences specific to one or few cell types in genes with expression enriched during development^19^. There remains an unmet need to uncover possible mechanisms of sex differences in autosomal gene expression in the prenatally developing cerebral cortex. In particular, studies of gene expression regulation could reveal noncoding cis-regulatory elements (CRE) harboring pathogenic variants, as well as inherited or *de novo* neurodevelopmental disorder (NDD)-associated variants^20,21^.

Studies of sex differences in the developing brain have focused on brain regions with enriched hormone signaling, such as the hypothalamus and amygdala^22^. Prenatal development represents an important period of testosterone production by gonadal cells of male individuals, which peaks around mid-gestation^23–25^. Both androgen (including testosterone) and estrogen signaling have been shown to regulate the development of the cerebral cortex^26,27^. Further, androgens and estrogens have been implicated in the control of autism risk gene expression^5^. Thus, hormone signaling could represent a major source of autosomal gene expression differences between males and females during prenatal development of the human cerebral cortex. However, we currently lack a systematic assessment of genomic targets of hormone receptors in the human cerebral cortex. Characterization of hormonal, as well as nonhormonal, contributions to sex differences during development represents a potential mechanism of sex-biased penetrance of autism.

Here, we address two main hypotheses that could explain the sex-biased prevalence of autism: first, that gene expression and regulation differences lead to greater vulnerability in males compared to females; and second, that sex differences might emerge as a consequence of hormone (androgen and estrogen) signaling, where differences in hormone levels during prenatal brain development may contribute to higher susceptibility of males to autism^28–30^. To this end, we applied a comprehensive genomic approach to identify molecular signatures of sex differences in the developing human cerebral cortex at midgestation, resolving the contribution of the X and Y chromosomes, hormones, or other epigenetic differences to sex-biased gene expression. By analyzing sex-specific burdens of *de novo* variants, we identify an overlap between sex differences and NDD genetic risk architecture to enable systematic and comprehensive studies of mechanisms that could contribute to male-biased diagnosis rates in autism.

## Results

### Integrated single-nucleus multimodal analysis of the midgestation brain

We applied single-nucleus Multiome (snMultiome) analysis to measure gene expression and chromatin accessibility in 21 female and 27 male midgestation human brain samples (**Fig. 1a**, **Fig. S1a, Supplementary table 1**), employing a pooling strategy designed to minimize technical batch effects. After removing low-quality cells, doublets, and cells with ambiguous sex, we retained 38,536 cells (19,209 female and 19,317 male) across 11 major cell types identified by enriched expression of known cell-type markers, including layer-specific excitatory neurons (ENs), medial and caudal ganglionic eminence derived inhibitory neurons (IN-MGE, IN-CGE, respectively), intermediate progenitor cells (IPCs), and radial glia (RG) (**Fig. 1b-c**, **Fig. S1b-e, Supplementary table 2**). We captured 227,118 open chromatin regions across all cell types (**Fig. 1d, Fig. S1f-g, Supplementary table 3**). To optimize genomic characterization of these regions, we further profiled putative promoter and enhancer histone marks (H3K4me3 and H3K27ac, respectively), as well as the repressive chromatin mark H3K27me3 and transcription factor CTCF by CUT&Tag^31^ for a subset of male (n = 6) and female (n = 5) samples included in our snMultiome analysis (**Fig. S1h, Supplementary table 4-7**). Integrating these epigenetic marks with the ATAC-seq data, we identified 17,157 promoters, 26,658 intergenic, 52,081 intronic, and 131,222 peaks at undefined genomic regions (**Supplementary table 3**). Integrating the RNA and ATAC assays allows for identification of enriched transcription factor motifs in accessible chromatin where the associated transcription factor is also expressed in the same cell, highlighting likely gene regulatory modules in a cell-type-specific manner (**Fig. 1e-f, Fig. S1i**).

**Fig. 1:**
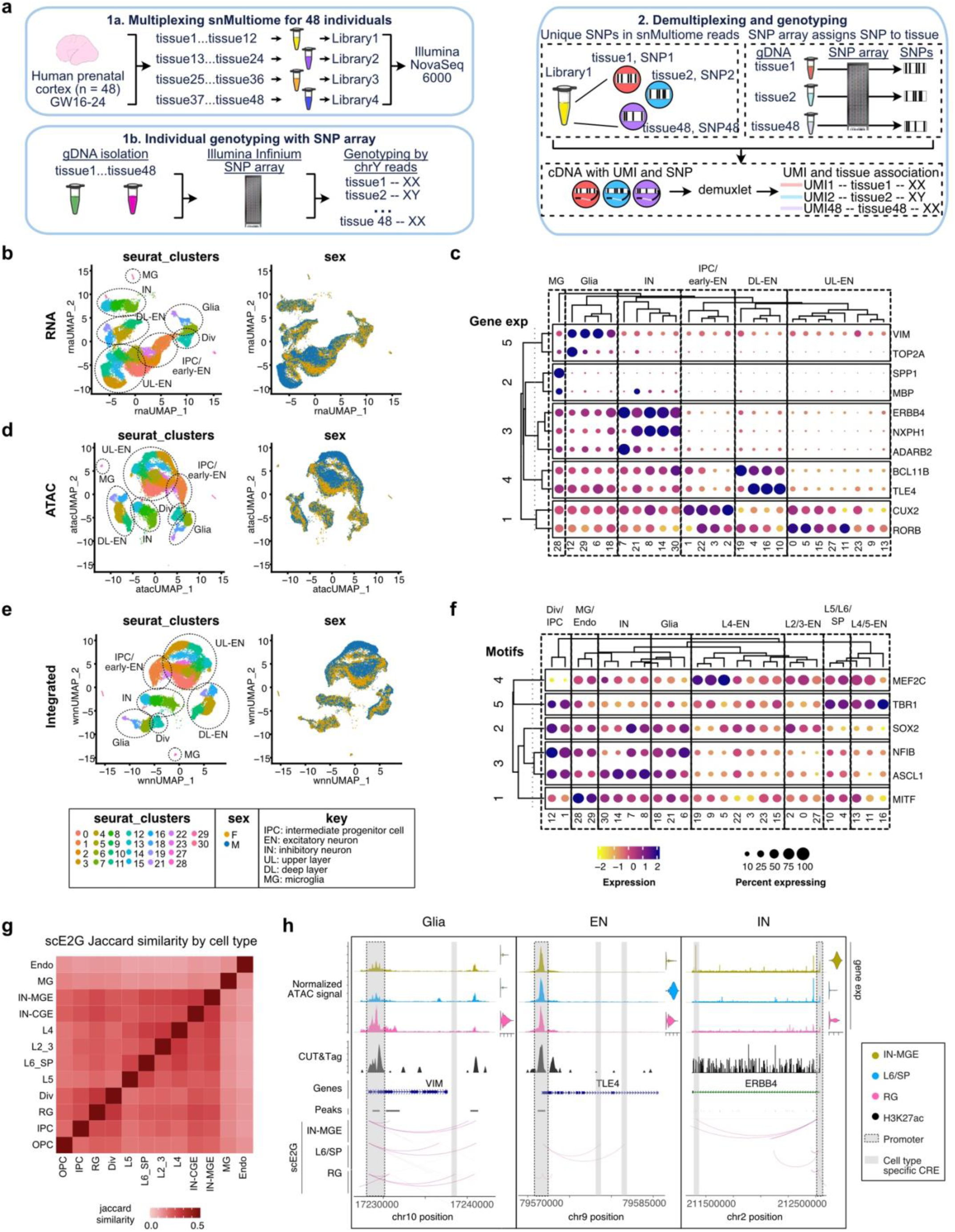
An atlas of male and female mid-gestation gene expression and chromatin accessibility. **a)** Schematic of the single cell Multiome assay (scMultiome) (1a) and single nucleotide polymorphism (SNP) array genotyping (1b) for 48 prenatal samples (21 female, 27 male). **b)** Schematic of donor genotype deconvolution (demuxing) of scMultiome data using SNP genotypes. **c)** UMAP of RNA assay by cell type and sex. **d)** UMAP of ATAC-seq assay by cell type and sex. **e)** UMAP of integrated assay by cell type and sex. **f)** Hierarchical dot plot of top gene expression cluster markers. **g)** Hierarchical dot plot of top enriched motifs by cluster. **h)** Correlation matrix of Jaccard similarity scores for CREs predicted for each cell type by the scE2G pipeline. **i)** Coverage plots showing cell type-specific CREs. Normalized ATAC-seq signal, gene expression, H3K27ac CUT&Tag, and scE2G predicted CREs by cell type are shown. Curved lines represent predicted enhancer-promoter interactions. To represent the three most distinct cell types, radial glia (glia) layer 6/subplate excitatory neurons (EN), and MGE derived inhibitory neurons (IN) are shown.

To identify potential regulators of gene expression, we predicted cell-type-specific functional enhancer-gene interactions using the single-cell Enhancer to Gene pipeline (scE2G)^32^ (**Supplementary tables 8-19**). As expected, transcriptomically similar cell types showed greater similarity among CREs (**Fig. 1g**). About 10-25% of predicted functional CREs are cell-type specific (**Fig. S1j**). As expected, these cell-type-specific CREs were predicted to regulate known cell type markers, such as *VIM* in radial glia (RG), *TLE4* in layer VI/subplate excitatory neurons (L6/SP), and *ERBB4* in MGE inhibitory neurons (IN-MGE) (**Fig. 1h**).

### Sex differences in gene expression

To identify sex differences in gene expression between males and females, we pseudobulked log-normalized RNA counts by cell type and used a linear mixed model to identify additional sources of gene expression variation (including batch and age, **Methods**)^33^. We identified 943 nonredundant differentially expressed genes by sex (DEGs, 730 female biased, 205 male biased, 8 with variable sex bias with cell type) in at least one cell type (FDR < 0.05, log2FC > |0.3|) (**Fig. 2a, Fig. S2a-b, Supplementary table 20**). Of the 943 total DEGs, 883 are autosomal genes (685 female-biased, 191 male-biased, 7 with variable sex bias with cell type), including the autism risk genes *FOXP2* (female biased, IN-MGE), and *MEF2C* (female biased, IPC) that have been previously reported to show female bias in the midgestation brain^19^ (**Fig. 2b, Supplementary table 20**). Out of 738 female-biased genes, 219 (29.7%) are differentially expressed across multiple cell types (**Fig. S2b**), of which 16 (7.3%) are encoded on the X chromosome, including the known X chromosome inactivation (XCI) escapees *DDX3X* and *ZFX*^34^, as well as genes predicted to be subject to XCI, such as *PDHA1* and *NAP1L3*. The other 203 genes with female-biased expression in multiple cell types are autosomal, including *H3F3A* and *MEF2C*. In contrast, 25 out of 213 (11.7%) of male-biased genes are differentially expressed in more than one cell type, with 9 out of 25 (36%) of these genes encoded on the Y chromosome (including *DDX3Y, KDM5D,* and *ZFY*) (**Fig. 2b, Fig. S2b**). With the exception of one lncRNA gene, the remaining eight out of nine male-biased Y-chromosome genes have an X-chromosome paralog, while only two genes out of 32 female-biased X genes have a chromosome Y paralog (**Supplementary table 20**). Both chromosome Y paralogs are male-biased in expression (*DDX3X-DDX3Y, ZFX-ZFY*) (**Fig. 2b**).

**Fig. 2:**
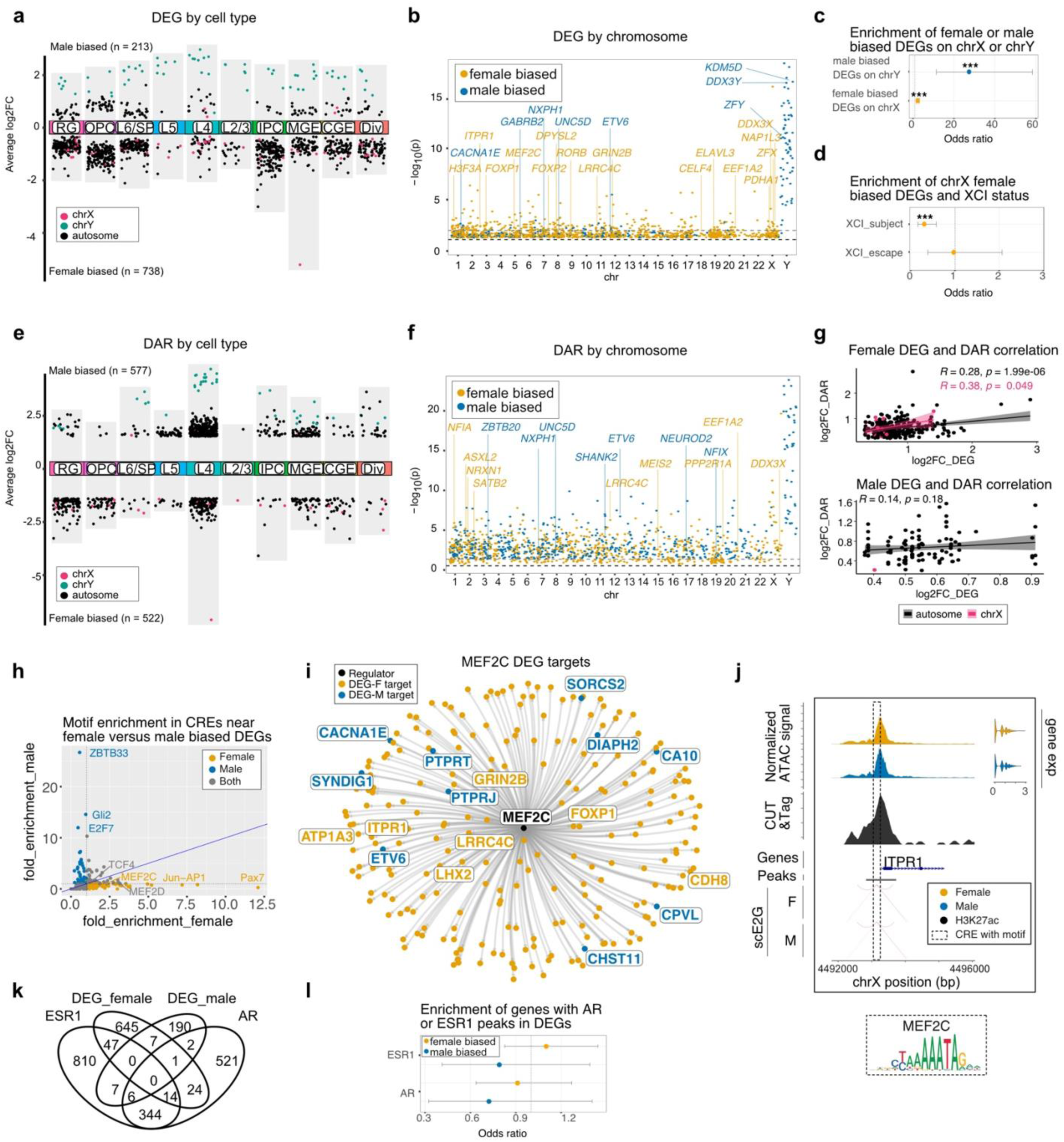
Sex-specific differences in gene expression and chromatin accessibility. **a)** Differentially expressed genes (DEGs) between male (XY, positive log_2_FC) and female (XX, negative log_2_FC) cells by cell type (FDR < 0.05). MG and Endo cell types were excluded due to cell counts <150. **b)** Distribution of DEGs by autosome, X and Y chromosomes, separated by sex-biased expression. Black dashed line indicates adjusted p = 0.05, grey dashed line indicates adjusted p = 0.01. **c)** Enrichment of female biased genes on the X chromosome, and male biased genes on the Y chromosome determined by a two-sided Fisher’s exact test. 95% confidence interval represented in grey. *** p < 0.001. **d)** Enrichment of genes either subject to or which escape from X chromosome inactivation (XCI) among X chromosome female-biased DEGs determined by a two-sided Fisher’s exact test. 95% confidence interval represented in grey. **e)** Differentially accessible regions (DARs) between male (XY, positive log_2_FC) and female (XX, negative log_2_FC) cells by cell type. Genes on autosomes, chrX and chrY are in black, pink, and green, respectively. FDR < 0.05, log_2_FC < |1.25|. MG and Endo cell types excluded due to cell count <150. **f)** Distribution of DARs by autosome, chrX, and chrY, separated by sex-biased expression. A subset of nearest genes predicted to be regulated by DARs are labeled. Black dashed line indicates adjusted p = 0.05, grey dashed line indicates adjusted p = 0.01. **g)** Scatter plot and Pearson correlation for genes that are both differentially expressed (X-axis) and have a DAR (Y-axis) by sex. Pearson correlation was calculated separately for genes on autosomes (black), X chromosome (pink), and Y chromosome (green). **h)** Scatter plot of motif enrichment for scE2G CREs for DEGs, with motif enrichment in CREs near female- biased DEGs on the x-axis, and motif enrichment in CREs near male-biased genes on the y-axis. Genes with fold enrichment > 1 in females and < 1 in males are highlighted in orange, and those with fold enrichment > 1 in males and < 1 in females in blue. **i)** Regulon plot for MEF2C predicted targets that are also DEGs. Targets that are female-biased DEGs are orange (n = 225), targets that are male-biased DEGs are blue (n = 7) of a total of 737 targets. **j)** Coverage plot of female biased gene *ITPR1*, with normalized ATAC-seq tracks and gene expression by sex, as well as H3K27ac CUT&Tag coverage and scE2G CRE predictions by sex. The MEF2C motif is highlighted in grey. **k)** Venn diagram of gene overlaps between male- and female-biased DEGs, nearest neighbor genes to AR CUT&Tag peaks, and nearest neighbor genes to ESR1 CUT&Tag peaks. **l)** Two-sided Fisher exact test enrichment of genes with either ESR1 (top) or AR (bottom) CUT&Tag peaks in either female (orange) or male (blue) DEGs.

Consistent with prior bulk tissue studies^18^, X and Y chromosome genes were highly enriched among sex-biased DEGs across most or all cell types (**Fig. S2b**), with significant enrichment of X chromosome linked genes among female biased DEGs (two-sided Fisher’s exact test, p = 2.9e-05, odds ratio (OR) = 2.05) and Y chromosome linked genes among male biased DEGs (two-sided Fisher’s exact test, p = 3.2e-10, OR = 27.26) (**Fig. 2c**). Among the female-biased chrX DEGs, we find a significant under-enrichment of genes predicted to be subject to X chromosome inactivation (XCI, determined by San Roman 2023^34^, two-sided Fisher’s exact test, p = 1.4e-4, OR = 0.31, **Fig. 2d**), as expected, with neither under nor over enrichment of X-chromosome genes predicted to escape XCI (two-sided Fisher’s exact test, p = 1, OR = 0.94). Taken together, the observed gene expression differences suggest both chromosome X, as well as autosomal, contributions to female-specific gene regulation, while male-biased gene expression is predominantly driven by Y chromosome genes across all cell types.

### Sex differences in chromatin accessibility do not predict differential gene expression

To identify male- and female-biased differentially accessible regions (DARs), we again applied a linear mixed model to account for covariates such as age and tissue pool. We found 522 female biased regions and 577 male biased regions (FDR < 0.05, log2FC > |1.25|) across all cell types (**Fig. 2e, Fig. S2c, Supplementary table 21**). Interestingly, most male biased peaks are found at distal regulatory regions, while most female-biased peaks are found within gene bodies (**Fig. S2d**). The most significant differences are found on the X and Y chromosomes, respectively (**Fig. 2f**).

To test whether differences in chromatin accessibility are a major driver of gene expression differences in the developing cortex, we leveraged scE2G on all female cells and all male cells separately (**Supplementary tables 22, 23**) to predict CRE-gene links that could be mapped onto our DARs (**Fig. S2e, Supplementary tables 24, 25**). We did not detect significant correlation between differential accessibility and gene expression in either females (Autosome: Pearson’s R = 0.28, p = 1.99e-06; X chromosome: Pearson’s R = 0.38, p = 0.048) or males (Autosome: Pearson’s R = 0.14, p = 0.178, **Fig. 2g**, **Fig. S2f**), though some sex-biased DEGs have corresponding DARs, such as female-biased *DDX3X* and male-biased *NXPH1* (**Fig. S2g**).

We then asked whether transcription factor (TF) motif enrichment analysis might uncover a link between sex-biased gene expression and chromatin accessibility, with TFs that are differentially expressed predicted to bind to DARs. Motifs of several TFs, including NEUROD1, BHLHA15, and NEUROG2, were enriched among male- as well as female-biased DARs, suggesting common mechanisms of sex-biased epigenomic features. Among male DARs, we also identified enrichment of NF1 and TCF21 motifs, while female DARs were enriched for YY1, MEF2C/D, and TFE3 (**Fig. S2h, Supplementary table 26**). However, when we restrict our analysis to motifs that are female-biased (enrichment in female-biased DARs > 1 with Benjamini-Hochberg (BH)^35^ p < 0.05, and enrichment in male-biased DARs < 1) and male-biased (enrichment in male-biased DARs > 1 with BH corrected p < 0.05, and enrichment in male-biased DARs < 1), we find only 12 male-biased motifs, including TBR1, and EOMES, as well as 12 female-biased motifs, including X-chromosome linked TFE3, as well as basic helix-loop-helix and SOX TFs. None of the significant 12 female biased or 12 male biased motifs are differentially expressed, but extending this analysis to nominally male-biased TF motifs without BH corrected p < 0.05, we find that 5 out of 89 (5.6%) male biased TFs are differentially expressed with female bias (*CUX2, FOXP1, RORA, ZBTB18,* and *ZFX*). Further, *ZFX*, as well as 3 additional male biased TF motifs (*GATA1, ZNF41,* and *ZNF711*) are on the X chromosome (4 out of 89 nominally male-biased motifs, 4.5%). Only 2 out of 76 (2.6%) nominally female-biased TF motifs are on the X chromosome and neither are sex-biased in expression, while an additional 2 TFs are female-biased in expression (*MEF2C* and *USF2*).

### In silico prediction of sex-biased gene regulatory networks

Next, we compared motifs enriched within CREs of DEGs, since DEGs and genes with DARs are not well correlated. CREs of male biased DEGs were enriched for 37 TF motifs (enrichment in CREs near male-biased DEGs > 1 and BH adjusted p < 0.05, enrichment in CREs near female-biased DEGs < 1), including ZBTB33 (X-linked), GLI2, and E2F7. *E2F7*, a TF involved in DNA damage repair^36^ and cell cycle progression^37^, is the only TF with male-biased motif enrichment that is also male-biased in expression. Three out of 37 (8.1%) TFs with male biased motif enrichment are on the X chromosome (*ELF4*, *ELK1, ZBTB33*), with X-linked TF *ZNF41* nominally male biased (with 61 total nominally male-biased TF motifs).

CREs of female-biased DEGs were nominally enriched for 109 motifs, with only 1 TF on the X chromosome (*GATA1*). Restricting our analysis to TFs with significant female-biased enrichment (BH adjusted p < 0.05), we identify 100 TF motifs, including PAX7, JUN, and MEF2C (**Fig. 2h, Fig. S2i, Supplementary table 27**). Interestingly, *MEF2C* is a well-known NDD risk gene^38–40^ and is the only TF with female-biased motif enrichment that shows female-biased expression, suggesting a gene regulatory network driven by MEF2C that is unique to females. To identify MEF2C targets, we used Single-cell Multiomic Enhancer-based Gene Regulatory Network Inference (scMEGA)^41^. MEF2C is predicted to have the most target genes that are also female-biased DEGs, some of which are further validated by the presence of a MEF2C motif at the promoter, as with *ITPR1* (n = 737 total targets, n = 222 female-biased targets, n = 10 male-biased targets, **Fig. 2i-j, Supplementary table 26**). Taken together, these findings suggest that female biased molecular features may in part be driven by the autosomal gene *MEF2C*.

While no transcription factors with male-biased motif enrichment are predicted to regulate DEGs, the male-biased DEG *LHX6* is predicted to have the most male-biased DEGs among its target genes (total target genes = 467, male-biased DEG targets = 27, female-biased DEG targets = 18, **Fig. S2j-k, Supplementary table 28**), highlighting its potential role in male-specific gene regulatory programs. However, LHX transcription factor (TF) family motifs, including LHX1, LHX2, LHX3, LHX9, are more significantly enriched in CREs near female-biased DEGs compared to those near male-biased DEGs. Interestingly, the inverse result is seen in DARs, with more significant enrichment in male-biased compared to female-biased DARs. While inconclusive, these results suggest that LHX6 might be responsible for male-biased gene regulatory networks, either through expression of *LHX6* itself, binding within male-biased DARs, or promoting expression of other male-biased genes.

### Estrogen and androgen receptor binding explain minimal sex differences in gene expression

To probe if differences in gene expression likely arise as a consequence of sex hormone regulation of gene expression, we employed CUT&Tag for estrogen receptor alpha (ERa, *ESR1*) and the androgen receptor (AR) in a subset of prenatal samples (5 males and 6 females, GW20) to identify possible genomic targets of steroid hormone signaling in the developing human cerebral cortex (**Fig. S2l**). We identified 2,179 ESR1 consensus peaks, and 1,161 total AR peaks (**Supplementary tables 29, 30**). To identify candidate target genes, we assigned ESR1 and AR bound peaks to genes by proximity using ChIPpeakAnno and overlapped candidate ESR1 and AR target genes with sex-biased DEGs. Surprisingly, only 6% of female-biased DEGs contained an ESR1 peak (**Fig. 2k-l**), and only 6% male-biased DEGs contained ESR1 peak, and 1% contained an AR peak. There were no differences in either ESR1 or AR peaks between male and female samples (**Fig. S2m**).

### Genes with sex-biased enrichment for *de novo* mutations

We sought to better understand the genetic risk architecture of autism in males and females, seeking to identify genes or pathways that might present males or females with greater vulnerability. Towards this goal we re-analyzed whole genome sequencing (WGS) and whole exome sequencing (WES) data from parent-child trios consisting of 61,743 NDD cases (30,687 autism and 31,056 DD) from the SPARK study (iWESv2) as well as five published cohorts of autism and developmental delay (DD)^2,3,42–44^ (**Fig. 3a, Supplementary table 31, Methods**). To predict genes that harbor a significant excess of *de novo* mutations (DNMs, including *de novo* likely gene-disruptive variants and *de novo* missense) in male probands (n=41,961) and female probands (n=19,782) separately (**Fig. 3a**), we applied recently described models of DNM enrichment that estimate the number of expected DNMs by incorporating locus-specific mutation rates, the gene length, and the null expectation based on chimpanzee–human coding sequence divergence (CH model)^3,45^, while the denovolyzer (DR) model estimates mutation rates based on trinucleotide context and adjusts divergence based on macaque–human gene comparisons^3,46^. 35 and 32 genes show the enrichment of DNMs specific to females and males respectively, henceforth referred to as female-biased and male-biased respectively (NDD_male32 and NDD_female35, **Fig. 3b, Supplementary table 31**). Male-biased genes were enriched for functional categories including transcription corepressor binding and transcription coregulatory binding, whereas female-biased genes were enriched for molecular functions such as histone modifying activity and N-acetyltransferase activity (**Fig. S3a**). Protein-protein interaction analysis identified a significant enrichment of interactions among all male-biased NDD genes, as well as all female-biased NDD genes, suggesting potential convergent molecular phenotypes with mutations in different sex-biased genes (**Fig. S3b**). After direct comparison of the DNM counts between males and females, only five genes (*DDX3X, HDAC8, WDR45, MECP2,* and *NAA10*) reached significance after correction for multiple comparisons (two-sided Fisher’s exact tests with BH correction) for an excess in females when compared to males, with another 20 genes reaching nominal significance (**Fig. 3b**, indicated with a single asterisk). By these criteria, no genes reached significance for excess of DNMs in male probands when compared to females, consistent with prior reports^3,47^. All 5 female-biased genes are encoded on X chromosome, consistent with the likely possibility that mutations in those genes are lethal in males due to the presence of a single copy of the X chromosome^3,48,49^ or lack of functional compensation from the Y-linked paralog^50^.

**Fig. 3:**
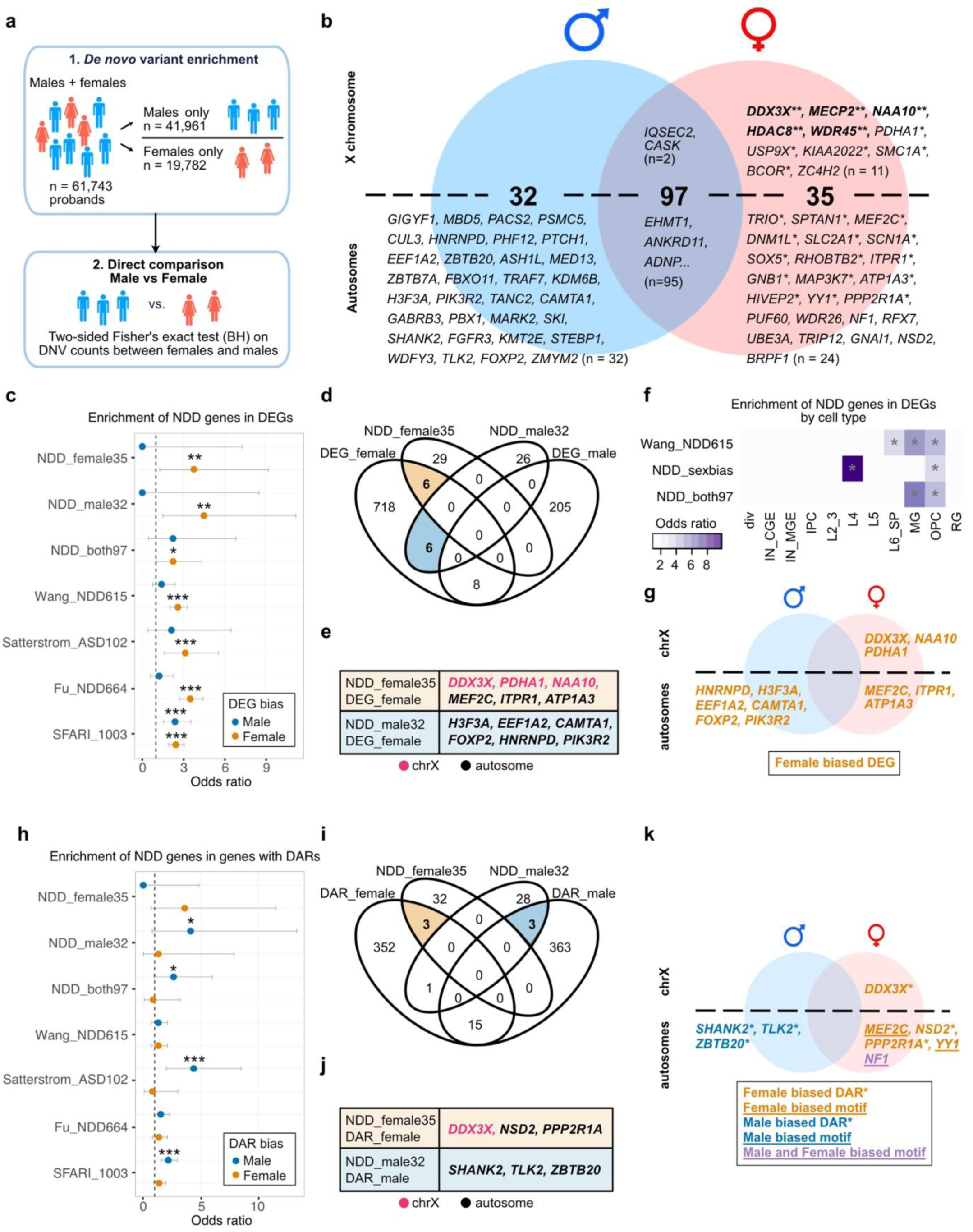
Gene expression and chromatin accessibility differences underlying NDD vulnerability. **a)** Schematic of sex-biased gene discovery. *de novo* mutation (DNM) enrichment analyses (1) were performed independently in NDD males and females using CH and DR models. Sex-biased genes were further identified by two-sided Fisher’s exact tests (BH correction, n=20,000 genes) of DNM counts between males and females (2). **b)** Venn diagram of genes with potential sex bias. Genes enriched for DNMs in males (n=129) and females (n=132) are shown in blue and red circles, respectively. Genes in bold with two asterisks (**) show sex-specific significance after multiple testing correction (a). Genes with a single asterisk (*) show nominal significance in the direct female-male comparison. **c)** Enrichment of NDD genes from different studies and DEGs, determined by two-sided Fisher’s exact test. Odds ratio shown in orange for enrichment in female biased DEGs and blue for male biased DEGs, with 95% confidence intervals shown in grey. *p < 0.05, **p < 0.01, ***p < 0.001. **d)** Venn diagram of differentially expressed genes and sex-biased NDD genes. **e)** Overlapping genes from NDD_female35 and DEG_female union (orange) and NDD_male32 and DEG_female union (blue). Overlapping genes on the X chromosome and autosomes are labeled in pink and black, respectively. **f)** Enrichment of NDD genes and differentially expressed genes (both male and female) by cell type. Wang_NDD615 contains all NDD genes identified in Wang 2022^3^. NDD_sexbias contains 35 female biased and 32 male biased genes from (B). Asterisk indicates p < 0.01 (Benjamini-Hochberg correction) by two-sided Fisher’s exact test. Color indicates odds ratio. **g)** Venn diagram from (b) highlighting the sex-biased NDD genes that are also differentially expressed. **h)** Enrichment of NDD genes from different studies and DARs, determined by two-sided Fisher’s exact test. Odds ratio shown in orange for enrichment in female biased genes with DARs and blue for male biased genes with DARs, with 95% confidence intervals shown in grey. *p < 0.05, **p < 0.01, ***p < 0.001. **i)** Venn diagram of nearest neighbor genes with DARs and sex-biased NDD genes. **j)** Overlapping genes from NDD_female35 and DAR_female union (orange) and NDD_male32 and DAR_male union (blue). Overlapping genes on the X chromosome and autosomes are labeled in pink and black, respectively. **k)** Venn diagram from (b) highlighting the sex-biased NDD genes that also either have a DAR or have sex- biased motif enrichment.

### Intersection of NDD genes and sex-biased gene expression

Next, we asked whether intersecting sex-biased molecular signatures of cortical development and NDD risk genes could illuminate the molecular mechanisms underlying sex bias of autism/NDD penetrance. To identify general patterns, we tested for enrichment of NDD genes within our entire DEG list agnostic of cell type, including several key NDD gene sets: NDD risk genes from Wang et al.^3^ (Wang_NDD615), NDD_male32 genes, NDD_female35 genes, NDD genes that enriched for DNMs in both sex groups (NDDboth97), 102 autism risk genes from Satterstrom et al.^1^ (Satterstrom_ASD102), NDD risk genes from Fu et al^2^ (Fu_NDD664), and SFARI genes with a score of 1 or 2 (SFARI_1003). Interestingly, both male and female biased NDD genes from this study, as well as genes without sex bias, show enrichment in female-biased DEGs (**Fig. 3c**). We identified significant enrichment of 6 male-biased autosomal NDD risk genes with female-biased expression: *H3F3A, EEF1A2, CAMTA1, FOXP2, HNRNPD, PIK3R2* (two-sided Fisher’s exact test, p = 4.2e-03, OR = 3.81) (**Fig. 3d-e**). This suggests that higher expression in females provides a buffer from potential damaging variants, where these variants would lead to an NDD diagnosis in males but not females.

Further, we identified significant enrichment of 6 female-biased NDD genes in female-biased DEGs: *DDX3X, PDHA1, NAA10, MEF2C, ITPR1, ATP1A3* (two-sided Fisher’s exact test, p = 0.02, OR = 2.77, **Fig. 3d-e**). Half of the candidate genes in this category are X-linked (*DDX3X, PDHA1, NAA10*), as expected, since two X chromosomes in females allow for up to twice the dosage of X-linked genes compared to males. For the female-biased X-linked NDD genes that are not female biased in expression, we hypothesize mosaicism in females where the wild type X chromosome is stochastically expressed due to random XCI, causing the X chromosome carrying the DNM to be incompletely penetrant. Therefore, baseline X-linked gene expression differences, as well as X chromosome mosaicism, likely contribute to sex-specific burden of *de novo* variation in these genes, where females can tolerate these mutations due to the increased gene dosage.

Additionally, we identified three autosomal (*MEF2C, ITPR1, ATP1A3*) female-biased NDD risk genes that are female-biased in expression. *ITPR1* and *ATP1A3* are both predicted transcriptional targets of MEF2C (**Fig. 2i**) that additionally have MEF2C motifs within regulatory regions (**Fig. 2j**). We therefore predict that higher expression of *MEF2C* is required for female-biased gene regulatory programs, and that this represents a point of NDD vulnerability biased towards female diagnosis. Additionally, we find NDD risk genes with no sex bias in *de novo* mutations that are both male and female biased in expression (**Fig. 3c**).

To assess any cell-type-specific vulnerability, we next intersected our DEG list by cell type with Wang_NDD615, as well as the sex-biased NDD genes in either males (32 genes) or females (35 genes) (NDD_sexbias), and the 97 NDD genes with no sex bias at the union of the Venn diagram (NDD_both97, **Fig. 3f, Fig. S3c**). Interestingly, DEGs in layer IV excitatory neurons show the greatest enrichment of NDD genes enriched in either males or females (adjusted p = 4.4e-03, OR = 9.82, one-sided Fisher’s exact test with BH correction), while there is no significant overlap with NDD genes that are enriched in both males and females (adjusted p = 0.341, OR = 2.44); out of 65 DEGs in this cell type, three are enriched in either male or female NDD probands, *DDX3X* (chr X), *PDHA1* (chr X), and *HNRNPD* (chr 4) (**Fig. 3g**). All three of these genes are more highly expressed in female individuals, but *DDX3X* and *PDHA1* are enriched in female probands, while *HNRNPD* is enriched in male probands (**Fig. 3b**). Previous work has demonstrated the convergence of co-expression networks related to autism within deep layer glutamatergic neurons^51^. These results further highlight that sex differences in gene expression related to NDD pathobiology are greatest in layer IV glutamatergic neurons. Overall, we find that genes with female biased expression, but not male-biased expression, are significantly enriched for both sex-biased and non-sex-biased NDD genes (**Fig. 3g**).

### Intersection of NDD genes and sex-biased gene regulation

We next applied this framework to differential gene regulation as a point of sex-biased vulnerability to NDD. We looked for an enrichment of NDD genes from the same datasets above (Wang_NDD615, NDD_male32, NDD_female35, NDD_both97, Satterstrom_ASD102, Fu_NDD664, and SFARI_1003) in genes with DARs, where the gene might not be expressed at the developmental time assayed, but may be differentially regulated between males and females at other points during development. Interestingly, we found that genes with male-biased DARs are enriched for NDD risk genes across multiple datasets, while there is no significant enrichment of female-biased DARs for these genes (**Fig. 3h**, two-sided Fisher’s exact test). This significant enrichment is seen for NDD_male32 in male-biased DARs (p = 0.044, OR = 4.10), with three genes that overlap: *SHANK2, TLK2,* and *ZBTB20* (**Fig. 3i-j**). While the enrichment of NDD_female35 in female-biased DARs is not significant (p = 0.06, OR = 3.59), there are three additional genes that have both a female-biased DAR and are also female-biased NDD risk genes: *DDX3X, NSD2*, and *PPP2R1A* (**Fig. 3i-j**). This suggests that sex-specific accessible chromatin, and therefore gene regulation, might underlie both male and female-biased vulnerability. There is no enrichment of NDD risk genes with no sex bias (NDD_both97) in genes with female-biased DARs, while genes with male-biased DARs are moderately but significantly enriched for these genes (**Fig. 3h, Extended data Fig. 3f**). Unlike our differential gene expression analysis, we find little cell-type-specific enrichment of NDD genes in genes with DARs (**Fig. S3g**).

To further assess whether gene regulatory differences might lead to differences in NDD vulnerability, we looked at female- and male-biased motifs (**Fig. 2h, Fig. S2h-i, Supplementary tables 25, 26**) to see if any TFs in our sex-biased NDD gene lists were also predicted to differentially regulate male and female biased genes. From our motif enrichment analysis in DARs, we identified NF1-halfsite motifs enriched in both female (p = 3.45e-5, fold enrichment = 1.1) and male (p = 1.32e-36, fold enrichment = 1.3) biased DARs, where NF1 is a female-biased NDD risk gene. Additionally, *YY1* and *MEF2D* (similar to *MEF2C*) are 2 of the top 10 most female-enriched motifs (**Fig. S2h**), where *YY1* and *MEF2C* are female-biased NDD risk genes. *MEF2A* and *MEF2B* are also enriched in female-biased DARs, but not male, further supporting the importance of the MEF2 TF family in female-specific gene regulatory programs, and the possibility that this might represent a point of female-biased NDD vulnerability. Interestingly, *YY1* is an autosomal gene with binding heavily biased towards the inactive X chromosome^52,53^, in addition to its role as a transcriptional activator^54^, highlighting how autosomal genes may regulate gene expression and chromatin state on the X chromosome. Lastly, though not a sex-biased NDD risk gene, *TFE3* is also an X-linked NDD risk gene identified in our Wang_NDD615 dataset, and one of the top 10 motifs enriched in female-biased DARs.

These findings highlight how sex-biased NDDs, while they may not be differentially expressed themselves, have the capacity to regulate sex-biased genes, and therefore lead to sex-specific effects with *de novo* variation. Additionally, the significant enrichment of NDD risk genes in genes with female-biased differential expression, but male-biased in accessibility, suggests that coding variation underlies female-specific protection for autosomal genes, but vulnerability on the X chromosome, while noncoding variation might underlie male-biased vulnerability. Previous work suggests that the regulatory burden of noncoding variation on the X chromosome specifically underlies male-biased vulnerability to NDD penetrance, where females are protected by the second copy of the X. We therefore sought to test the regulatory burden of inherited variation in both males and females, with a focus on the X chromosome.

### The X chromosome contributes to NDD vulnerability through sex-specific mechanisms

To further interrogate the contribution of the X chromosome to sex-biased NDD vulnerability, we first established an integrative map of genes that are differentially expressed, have sex differential accessibility, and confer NDD risk via our DNM analysis (**Fig. 4a)**. We additionally mapped our CREs identified in bulk female samples, as well as bulk male samples. Since chromatin accessibility does not correlate with differences in expression, most DEGs on the X chromosome do not have an additional DAR, with the exception of *EIF1AX* and *DDX3X*. Of the 51 chrX DEGs, 7 are NDD risk genes: *CDKL5, CNKSR2, NONO, DCX, DDX3X, NAA10*, and *PDHA1*. 5 chrX DEGs are male-biased: *AC233296.1, CNKSR2, CLIC2, PLS3*, and *REPS2*. There are 18 total unique DARs, 15 with female bias and 3 with male bias.

**Fig. 4:**
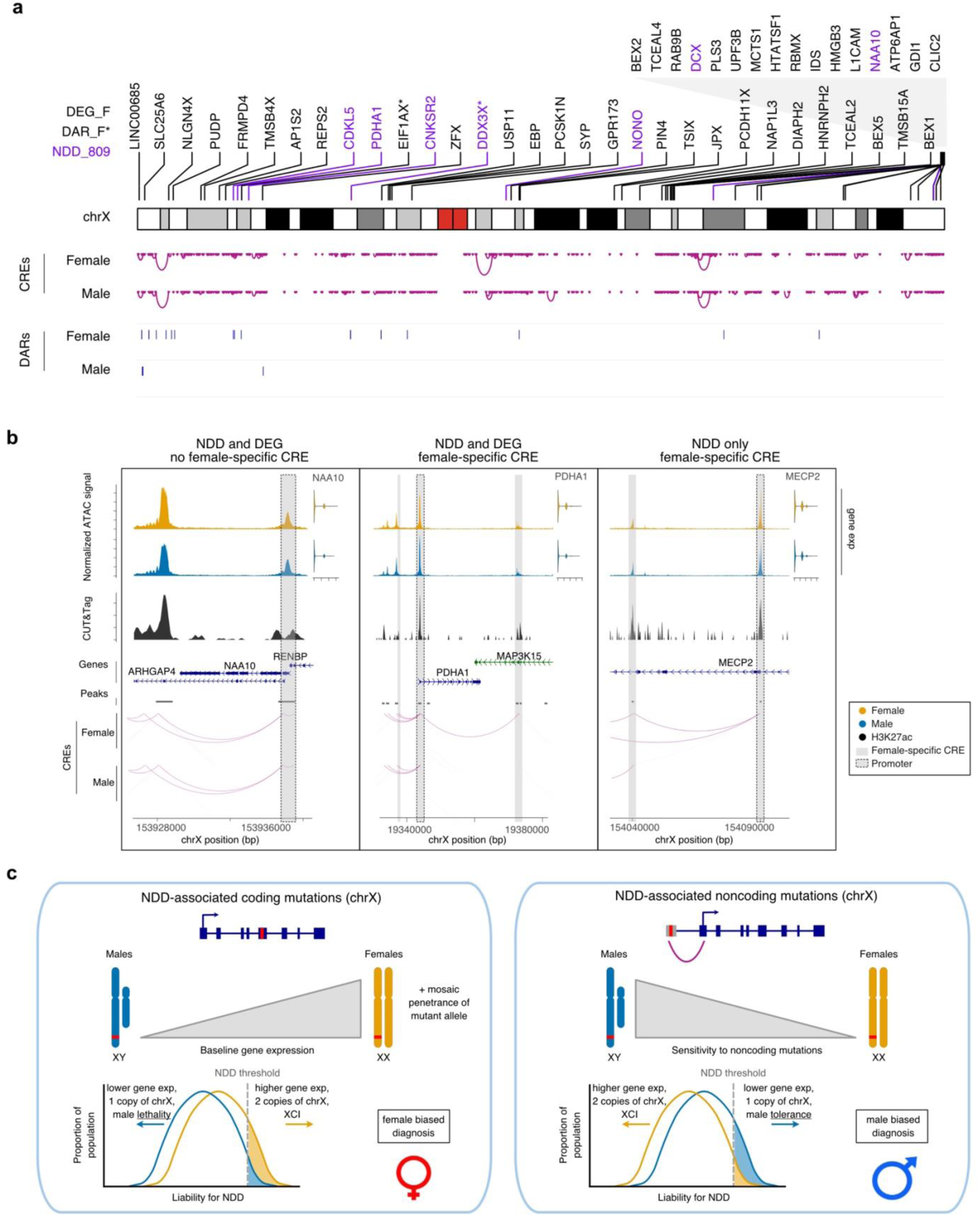
An integrated genomic map of the X chromosome. **a)** Map of chromosome X including DEGs, DARs, NDD genes that are also differentially expressed, and CREs. **b)** Three examples of chromosome X NDD genes: *NAA10*, *PDHA1*, and *MECP2*. **c)** Proposed model of sex-specific vulnerability. X chromosome coding mutations lead to female-biased vulnerability, and noncoding mutations lead to male-biased vulnerability.

While about 70% CREs identified in both female and male bulk analyses do not show any differences (48,315 shared CREs, 68,066 total bulk female CREs, 68,509 total bulk male CREs), as with CREs predicted to regulate female-biased DEG *NAA10*, about 30% of CREs are sex-specific (19,751 female-specific, 20,194 male-specific **Fig. 4b, Supplementary tables 22, 23**). For example, female-biased NDD risk genes *PDHA1* and *MECP2* have CREs predicted to interact with the promoter in females but not males.

Together, this comprehensive genomic map inclusive of the X chromosome highlights potential sources for sex-biased NDD vulnerability, including genes with female-biased expression that allow for tolerance to coding variation, as well as CREs that may carry noncoding mutational burden biased towards males who lack mosaicism and compensation from the wild type X chromosome (**Fig. 4c**).

## Discussion

In the current study, we identify cell-type-specific transcriptomic and epigenomic sex differences in the developing human cerebral cortex at midgestation. In addition to the genes encoded on the X and Y chromosomes, we identify dozens of autosomal genes with female-biased expression. Consistent with a prior analysis, we identify significant enrichment of high-confidence NDD risk genes among female-biased genes^19^. Autosomal gene expression differences detected in this study highlight the unique features of prenatally developing human brain in contrast to the adult brain, where transcriptomic differences seem to be largely restricted to X and Y chromosome linked genes^55^. Interestingly, the same extent of bias among autosomal signals was not found among chromatin accessibility signatures, which were more balanced between males and females.

The prenatally developing human brain is exposed to circulating hormones, including testosterone, whose levels surge during second trimester^23–25^, and which can be converted to estrogen. Hormone signaling plays critical roles in development of brain regions relevant to social interactions^56^, but recent studies have shown that these hormones can induce changes in gene expression and progenitor cell behavior in the cerebral cortex^26,27,57^. Our study did not uncover a strong correlation between ESR1 or AR binding sites and sex biased genes within the prenatally developing cerebral cortex, but binding of other sex hormone receptors including ESR2 and progesterone receptor, as well as inter-individual variability and earlier stages of development will need to be examined in the future. Analysis of developmental regulomes implicated transcription factor *MEF2C* as a major regulator of sex-biased expression of hundreds of autosomal genes in females. Similarly, our analysis implicates X-linked transcription factor TFE3 as a candidate driver of chromatin accessibility differences.

By intersecting the map of molecular sex differences with genetic risk architectures of male and female individuals with autism, our study presents a resource for exploring the molecular basis of sex-biased cortical development and autism diagnosis. We highlight the role of the X chromosome as a driver of molecular sex differences in the developing cortex. The expected lethality of high penetrance X-linked NDD risk genes in males results in strong bias towards the diagnosis of coding mutations in these genes in females with autism and NDDs^58^. However, emerging evidence from studies of mutations disrupting the activity of CREs of X-linked NDD risk genes offers clues into a mechanism by which X chromosome mutations could result in autism in some males because of their smaller genetic impact on gene expression^59^. This finding is supported by recent studies implicating noncoding elements in NDD risk^21^, as well as male-specific burden of inherited variation on the X chromosome driven by female protection of two copies of the X chromosome^59^.

The majority of autism and NDD studies have overwhelmingly focused on identifying genes with an excess of protein-damaging variants^2,3^, and the contribution of noncoding mutations to autism remains underexplored. Our resource defines a set of CREs of X-linked NDD risk genes that could be used to prioritize noncoding variants for functional validation. Thus, a shift from generating familial whole-exome sequencing data to whole-genome sequencing data will be critical to testing genetic burden among CREs of NDD risk genes. The larger whole-genome sequencing datasets being generated for autism families as part of SPARK^60^ in particular (currently 142,357 individuals from 54,558 families in SPARK iWESv3 and 12,519 individuals from 3,394 families in SPARK iWGSv1.1, https://base.sfari.org/public-resources) will allow us to test whether DNMs in CREs located on the X chromosome and other sex-biased genes contribute to the increased male-biased diagnosis of autism. The transcriptomic and epigenomic data resource presented here provides a refined annotation of genomic loci likely to be functional in the critical brain region and period of development most highly associated with autism^7,61^.

## Methods

### Tissue source

De-identified tissue samples were collected with previous consent in strict observance of the legal and institutional ethical regulations. Protocols were approved by the Human Gamete, Embryo, and Stem Cell Research Committee (institutional review board) at the University of California, San Francisco.

### Flash freezing tissue

Isopentane was cooled with liquid nitrogen, and the tissue was submerged in the isopentane for 30 seconds. Flash frozen tissue was transferred to cryovials and stored at −80C.

### DNA isolation and SNP genotyping

The DNA from each sample was isolated using an adapted protocol from the Purelink Genomic DNA Kits (Fisher K182001). 15 mg of flash frozen tissue samples were shaved off from tissue blocks on dry ice using pre-chilled 60x15 mm plates and sterile blades. The shaved tissue was transported to regular ice and 210 uL of master Digestion Buffer containing 180 uL of Purelink Genomic Digestion Buffer and 30 uL of Proteinase K was added to the sample. The combined sample and master Digestion Buffer was lysed mechanically using a chilled blade and transferred to a DNA Lobind tube (Eppendorf) using a wide-bore pipette tip. The sample was incubated at 55C for 4 hours at 600 rpm with vortexing every half hour to hour. The DNA was bound to the spin column, washed, and eluted using the Purelink Genomic DNA Kit (Fisher K182001). The isolated DNA was sent to UC Berkeley (QB3 Genomics, UC Berkeley, Berkeley, CA, RRID:SCR_022170) for SNP Genotype sequencing using the Illumina Infinium Global Screening Array on the iScan system. Genotype (XX vs XY) for each sample was determined using Illumina’s Genome Studio software.

### Nuclei isolation

Flash frozen tissue blocks were cut in groups of twelve on dry ice using prechilled 60x15 mm plates and sterile blades. Samples were pooled, with 12 individuals per nuclei isolation. For each of the twelve tissue samples used, 5mg was obtained using this method and added to a shared tube of frozen tissue. The shared tube was weighed after the addition of each sample to ensure equal input and individual representation.

The tube containing 60mg of pooled frozen tissue was emptied into a 2 mL glass dounce that had been pre-chilled on ice. 1 mL of homogenization buffer containing 0.2653 M Sucrose, 26.6 mM KCl, 5.31 mM MgCl2, 21.2 mM Tricine-KOH pH 7.8, Ultrapure H2O, 1 mM DTT, 0.5 mM Spermidine, 0.15 mM Spermine, 0.3% NP40, 1X Protease Inhibitor (Sigma #11873580001), and 0.6 U/μL RNAse Inhibitor was added to the dounce. The tissue was homogenized using 12 strokes from the dounce pestle A and 16 strokes from the dounce pestle B. The resulting homogenized solution was filtered through the cap of a pre-chilled sterile 5 mL Falcon round bottom tube (Corning 08-771-23) and centrifuged at 400g for 5 min at 4C in a fixed-angle centrifuge. The supernatant was removed and discarded. The remaining pellet was resuspended using 400μL of the homogenization buffer and mixed in with 1 volume of a 50% iodixanol solution, resulting in a 25% iodixanol layer. A 600μL volume of 30% iodixanol made from the 50% Iodixanol stock solution and homogenization buffer was slowly layered below the 25% layer. A 600μL volume of 40% Iodixanol solution made from the 50% Iodixanol stock solution and homogenization buffer was slowly layered below the 30% layer. The layered solution was centrifuged in a pre-chilled swinging bucket centrifuge for 20 min at 4C at 3,000g with acceleration level at 1 and deceleration level at 0. The iodixanol gradient solution post-centrifugation contained a distinct nuclei band located within the 30% iodixanol layer. The 25% layer was removed and discarded using a 200μL pipette. 200μL of the nuclei band was slowly removed and placed in a pre-chilled 1.5 mL DNA lobind tube (Eppendorf 022431021). The 200μL band collected from the gradient solution was diluted with 200μL of wash buffer containing 10mM Tris-HCL pH 7.4, 10mM NaCl, 3mM MgCl2, 1% BSA, 0.1% Tween-20, 1 mM DTT, 0.6 U/μL RNAse Inhibitor, and ultrapure H2O. The diluted nuclei solution was counted manually using a disposable hemocytometer (Bulldog Bio DHC-N51) and Trypan Blue. The solution was then spun down using a pre-chilled fixed angle centrifuge at 500g for 5 min at 4C. The supernatant was removed and discarded. The remaining nuclei pellet was resuspended using variable volumes of Diluted Nuclei Buffer containing 1x Nuclei Buffer (10x Genomics PN-2000207), 1 mM DTT, 1 U/μL RNase Inhibitor, Nuclease-Free H2O taken from Chromium Next GEM Single Cell Multiome ATAC + Gene Expression Kit (10x Genomics PN-1000283). The final concentration of the nuclei suspension was between 8,000 and 10,000 nuclei/μL to load into the 10x Multiome protocol.

### Single-nucleus Multiome

Each nuclei extraction of pooled donors was split into four lanes of Multiome (10x Genomics Chromium Next GEM Single Cell Multiome ATAC + Gene Expression PN-1000283) with 18,000 nuclei loaded per lane, with a targeted nuclei recovery of 12,000 nuclei per lane. Libraries were pooled to 4nM and sequenced on a Novaseq 6000 at a final concentration of 150pM, with targeted read depth of 20,000 reads per cell.

### snMultiome QC

Both scRNA-seq and scATAC-seq were preprocessed following the Seurat Weighted Nearest Neighbor (WNN) analysis for 10x Multiome. Briefly, sc-RNA-seq data was subsetted to include cells with between 1000 and 25000 UMIs, between 1000 and 10000 genes, and less than 10% mitochondrial reads. Data was normalized using SCTransform. sc-ATAC-seq data was subset to include cells with between 5000 and 70000 fragments.

Data was batch corrected using Harmony for the RNA and ATAC assays separately, UMAPs for each assay were generated, and scRNA- and scATAC-seq data was integrated using the command FindMultimodalNeighbors. The WNN UMAP was generated, and clusters were identified from the RNA assay with 0.8 resolution. Cell types were manually annotated using a reference dataset. Low-quality clusters were removed, as well as doublets and cells with ambiguous genotype (i.e., cells with XX genotype determined by Genome Studio as described above, but with expression of SRY, a Y chromosome gene).

### Demultiplexing individuals using SNPs

We used the SNP genotypes in each sample to deconvolute sample information, assigning each cell to its donor. To reduce noise, we first subdivided the snMultiome data to include only donors from each batch, then input these barcodes into Popscle (https://github.com/statgen/popscle) and subsequently Popscl Demuxlet^62^ (https://github.com/statgen/demuxlet) to assign each cell to a specific donor included in that batch. Genotype aware parameters and including intronic sequences were used. The Demuxlet output was then merged with Seurat Object using a common cell barcode from cellranger to assign a donor ID to each unique cell.

### Generation of cortex ATAC map

We supplemented the snATAC-seq data generated here with previously published snATAC-seq data to generate a map of chromatin accessibility in the fetal cortex (Ziffra 2021^11^). Using the Sinto tool (https://timoast.github.io/sinto/), in conjunction with dataset-specific cell-type and donor labels, we sex and cell-type pseudobulked. For the Ziffra datasets, we utilized the following cell types: early excitatory neuron, upper-layer excitatory neuron, deep-layer excitatory neuron, CGE-derived interneuron, MGE-derived interneuron, oligodendrocyte progenitor cell, IPC, and radial glia. For the Multiome dataset from this study, we used the following cell types: layer 2/3 excitatory neuron, layer 4 excitatory neuron, layer 5 excitatory neuron, layer 6/subplate excitatory neuron, CGE-derived interneuron, MGE-derived interneuron, oligodendrocyte progenitor cell, IPC, radial glia and dividing radial glia. Peak-calling was performed on each pseudobulk using MACS2 with the following parameters: --SPMR -q 0.01 -gs --nomodel --ext-size 50 --shift 100^63,64^. Using an index script^65^, we generated a set of consensus peaks from the pseudobulked MACS2 outputs and reads from the pseudobulked datasets over the peaks were counted using the featureCounts read quantifier as part of the Subread package^66^ (https://github.com/ShiLab-Bioinformatics/subread). The consensus peaks were filtered for peaks on chromosomes 1-22, X and Y and mean FPKM above 2.

### Cell type- and sex-specific peak calling and scE2G CRE predictions

Cell type- and sex-specific candidate enhancers were identified and linked to their predicted target genes within our snMultiome data using the single-cell enhancer-to-gene (scE2G)^32^ method’s standard multiome pipeline (https://github.com/EngreitzLab/scE2G). Pseudobulk fragment input files were generated for each analysis group using the SplitFragments function from the Signac v1.14.0 package^67^. Fragment files were sorted by coordinate using sortBed from the bedtools v2.27.1 package^68^ with the ‘-i’ flag. Samtools v1.21^69^ was then used to individually zip each fragment file using the bgzip command then index each zipped file using the tabix command with the ‘-p bed’ flag. RNA count matrix input files were generated by extracting raw counts for each analysis group from the Seurat object, ensuring that the cell IDs matched those which were present in the corresponding fragment files, then exporting each matrix to csv and compressing the resulting files using gzip. Pipeline configuration files were generated to follow the standard multiome prediction workflow using the provided ‘multiome_powerlaw_v3’ model. Once all files were prepared, the pipeline was run using the following command: snakemake -j1 --use-conda --configfile config/config.yaml. A threshold of 0.177 was used to generate the final CRE maps. Due to lack of power, we were not able to predict CREs on the Y chromosome.

### CUT&Tag

CUT&Tag was performed on flash frozen tissue as previously described^31^ with some modifications. 3mg of tissue was thawed and minced, then resuspended in PBS with 0.1% formaldehyde for 1 minute. 1.25M glycine was added to double the molar concentration of formaldehyde and stop cross linking, and cells were spun down at 1300g at 4C for 3 minutes. Tissue was resuspended in 1ml PBS and centrifuged at 2000g for 5 min at 4C. Tissue was then resuspended in wash buffer (20mM HEPES pH 7.5, 150mM NaCl, 0.5mM spermidine, and 1 EDTA-free complete protease inhibitor tablet) and homogenized using a glass dounce. Cells were transferred to a clean 1.5ml tube and centrifuged at 3000g for 3min at RT, and resuspended in 1ml wash buffer. Wash step was repeated twice more for a total of 3 washes. Concanavalin A-coated beads (Fisher Scientific #NC1526856) were prepared by adding 10µL beads per reaction to bead-binding buffer (20mM HEPES pH 7.9, 10mM KCl, 1mM CaCl2, and 1mM MnCl2). Using a magnetic rack, binding buffer was removed, and beads were resuspended in binding buffer once more before removing the binding buffer again and finally resuspending in enough binding buffer for 10µL per reaction. 10µL beads were then added to cells and incubated on an end over end rotator for 10 minutes at room temperature. Wash buffer was removed from cells using a magnetic rack, and cells were resuspended in enough antibody buffer (wash buffer with 2mM EDTA, 0.1% BSA, and 0.05% digitonin) for 50µL per reaction. Primary antibodies (H3K27me3 Cell Signaling Technologies #9733, H3K4me3 Cell Signaling Technologies #9751T, H3K27ac Cell Signaling Technologies #8173, CTCF Epicypher #13-2014, IgG Epicypher #13- 0042) were added at 1:50 dilution and samples were nutated overnight at 4C. Using a magnetic rack, primary antibody was removed, and cells were resuspended in 100µL secondary antibody (Antibodies Online #ABIN101961) diluted in wash buffer with 0.05% digitonin. Cells were then nutated for one hour at room temperature. Secondary antibody mix was removed using a magnetic rack, and cells were washed with wash buffer with 0.05% digitonin three times. pA-Tn5 preloaded with Nextera adapters (Epicypher #15-1117) was diluted 1:20 in dig-300 buffer (20mM HEPES pH7.5, 300mM NaCl, 0.5mM spermidine, 0.015% digitonin with 1 EDTA-free complete protease inhibitor tablet) and cells were resuspended in 50µL of pA-Tn5 mix. Cells were nutated for one hour at room temperature. Using a magnetic rack, pA-Tn5 mix was removed, and cells were washed three times in dig-wash buffer. Cells were resuspended in 300µL of tagmentation buffer (dig-300 buffer with 10mM MgCl2) and incubated in a 37C water bath for one hour. Cells were released from beads with addition of 10µL 0.5M EDTA, 3µL 10% SDS, and 2.5µL 20mg/ml proteinase K. Samples were vortexed and incubated in a heat block at 55C for one hour. Fragments were purified by adding 300µL phenol:chloroform:isoamyl alcohol (25:24:1 v/v) and sample to a phase lock tube (Qiagen #129046), and spun down at 16,000g for 3 minutes at RT. 300µL chloroform was added to each sample and spun down once more at 16,000g for 3 minutes at RT. The aqueous layer was added to a 1.5ml lo-bind tube with 750µL 100% ethanol and mixed by pipetting. Samples were cooled on ice and spun down at 16,000g for 15 minutes at 4C. Supernatant was decanted, and samples were washed once more with 1ml 100% ethanol and spun down at 16,000g for 1 minute at 4C. Ethanol was decanted, the pellet was air dried, and resuspended in 22µL water. Libraries were amplified by mixing 21µL sample, 2µL each i5 and i7 primers (Illumina #FC-131-2001), and 25µL NEBNext High Fidelity 2X Master Mix (NEB #M0541S) and using the following PCR cycle settings: 72C for 5 minutes, 98C for 30 seconds, 98C for 10 seconds, 61C for 10 seconds, repeat steps 3-4 15x, and a final 72C incubation for 1 minute. Libraries were purified using SPRI Select Reagent (Beckman Coulter #B23317) and eluted in 20µL water. Libraries were pooled to 2nM and diluted to a final concentration of 750pM with 2% PhiX spike in for sequencing using the Illumina NextSeq 2000 with a targeted read depth of ∼10 million reads per sample.

### CUT&Tag analysis

Reads were trimmed using TrimGalore, aligned to reference genomes for hg38 and E. coli using bowtie2 with parameters “--end-to-end --very-sensitive --no-mixed --no-discordant --phred33 -I 10 -X 700.” Duplicates were marked and removed with picard. Peaks for each sample were called using MACS2 with respective IgG controls for each sample with parameters “-q 0.01 -B --SPMR --keep-dup all --nomodel.” Consensus peak files were generated by concatenating bed files from all samples for each histone mark.

### Differential gene expression and accessibility analysis

Differentially expressed genes were identified using Dreamlet^33^ (https://github.com/GabrielHoffman/dreamlet). Briefly, ribosomal genes were removed, outliers were removed, and cells were pseudobulked by cell type using the aggregateToPseudobulk command with default parameters. Variance partition was run to identify variance due to genotype, batch, and age. Differential analysis was run to identify expression differences due to genotype. Genes with FDR < 0.05 were used for downstream analysis. For DARs, thresholds of FDR < 0.05 and log2FC > |1.25| were used.

### DAR and DEG correlation

We leveraged our CRE-gene links identified by scE2G (methods above) for either bulk XX samples or bulk XY to link DARs to target genes. Briefly, the scE2G bedpe file with CRE-gene links was intersected with the respective DAR bed file (bulk XY scE2G with male-biased DAR, and bulk XX scE2G with female-biased DAR). We then mapped the associated gene from the scE2G file onto the DAR file to identify target genes for DARs. Then, Pearson’s correlation was used to determine if there was a significant correlation between the log2FC enrichment of accessible chromatin (DARs) and gene expression (DEG) for each male and female.

### Motif enrichment analysis

Bed files were used as input for Homer (https://github.com/javrodriguez/HOMER), using the findMotifsGenome command with default parameters. The knownResults predictions were used, and fold enrichment for each motif was calculated by dividing the target enrichment by the background enrichment. Sex-specific motifs were determined by calling motifs with enrichment in one sex > 1 and enrichment in the other sex < 1.

### scMEGA gene regulatory network prediction

Gene regulatory network analysis was done using Single-cell Multiomic Enhancer-based Gene regulAtory network inference (scMEGA, https://github.com/CostaLab/scMEGA)^41^. Using our preprocessed snMultiome object, we first determined pseudotime trajectory with ArchR^70^ for computing TF binding activity with TF expression along this trajectory. Next, we created the motif slot of our ATAC assay in our snMultiome object by running getMatrixSet (using JASPAR2020 database), createMotifMatrix with hg38, and createMotifObject to be integrated into the metadata in the ATAC assay, where we subsequently used RunChromVar with BSgenome.Hsapiens.UCSC.hg38. To associate TF binding motifs with target genes, we linked peaks to genes with SelectGenes and correlated TFs with target genes by running GetTFGeneCorrelation using the chromVar TF assay, RNA assay, and inferred pseudotime trajectory. We lastly associated TFs with genes by running motif.matching, and finally inferred gene regulatory networks with GetGRN using our matched motifs, TF-gene correlations, and peak-gene links. The output is a dataframe with TF regulator, gene target, and correlation score, where we used TF-gene predictions with a score of 0.4 or greater.

### Genetic analysis of NDD cohorts and sex-biased gene discovery

Variants from the SPARK cohort were called by GATK^71^ (v3.7-0-gcfedb6) and FreeBayes^72^ (v1.1.0-3-g961e5f3) based on GRCh38. DNMs were discovered using developed scripts as described in Wang et al. 2022. Briefly, Candidate DNVs were defined as sites where sequencing depth greater than 9 in all trio members, and both parents were "0/0" with no alternate reads in parents (AD=",0"), while the child was "0/1" or "1/1", and child allelic balance >0.25, child genotype quality >20 (GATK) or sum of quality of the alternate observations >20 (FreeBayes), and supported by both GATK and FreeBayes call sets. DNVs in low complexity and recent repeat ^73^ and in outliers (samples with 8 DNMs, Poisson distribution) were filtered. The coordinates of DNMs were then converted to GRCh37 using UCSC LiftOver Tool^74^. We annotated *de novo* likely gene-disruptive (dnLGD, including frameshift, stop-gain, splice-donor and splice-acceptor) and *de novo* missense (dnMIS) variants by using Ensembl Variant Effect Predictor (VEP, v110.1)^75^ and CADD score (v1.3)^76^. DNMs with allele frequency > 0.1% in gnomAD exomes v2.1.1^77^ were filtered. Finally, we integrated DNMs from known females and males including samples from SPARKiWESv2^3^, SPARK pilot^42^, MSSNG and AGRE cohorts^43^, ASC and SSC cohorts^2^, and DD samples^44^. DNMs that were observed in both affected and unaffected (n=9,754) were removed.

CH^45^ and DR^46^ models were used to identify genes with a significant excess of DNMs in 41,961 NDD males and 19,782 NDD females separately3. Genes were considered significantly enriched if they harbored ≥2 DNMs and passed the 5% family-wise error rate threshold (p < 5.1E-07, five tests) in both models. Total DNV counts for each enriched gene were compared between males and females using two-sided Fisher’s exact tests, with p values adjusted for multiple testing using the Benjamini–Hochberg method35 (n=20,000 genes). DNM counts in X-chromosomal genes were normalized by the number of X chromosomes represented in the analyzed subset of females (n = 35,076) and males (n = 31,675). This analysis was also performed for DD (17,421 males and 13,635 females) and ASD groups (24,540 ASD males and 6,147 ASD females, and 14,254 and 7806 copies of X chromosome for males and females, respectively) and the results were listed in **SI table 30**.

### GO and PPI analysis of sex-biased NDD genes

Gene ontology analysis was done using the clusterProfiler package with command compareCluster using NDD_male32 genes and NDD_female35 genes as input, with molecular function ontology assessed. STRING v12.0 database^78^ was used to perform PPI network analysis. We used the multiple protein input option with default settings including "confidence for meaning of network edges", "textmining", "experiments", "databases", "co-expression", "neighborhood", "Gene Fusion", and "co-occurence" for active interaction sources", and "minimum required interaction score" of 0.400. Disconnected nodes were hidden from the network output but not from the enrichment analyses.

### DEG/DAR and NDD enrichment

We defined the following variables: q = number of genes that overlap, m = number of NDD genes, n = number of total genes expressed (15,000), k = number of DEG or genes with DAR. We used a 2x2 matrix defined as (n-k, m-q, k-q, q) to be used for two-sided Fisher exact test, unless otherwise stated. * p < 0.05, ** p < 0.01, *** p < 0.001.

## Data availability

Raw data are available at dbGaP. Processed data are available on the UCSC cell browser at https://cells.ucsc.edu/?ds=ssd-prenatal-cortex+rna.

## Code availability

All scripts used for data analysis are available at https://github.com/NOW-Lab.

## Acknowledgments

We thank all of the individuals who participated in this research. This work was supported, in part, by the Simons Foundation (SFARI #810018 to T.J.N., E.E.E., H.Y.Z., and A.C.), the US National Institutes of Health (NIH R01MH101221 to E.E.E.; R01NS123263 and R01MH125516 to T.J.N.), and NSF 2134955 to T.J.N., as well as gifts from the Esther A. & Joseph Klingenstein Fund, Shurl and Kay Curci Foundation, Sontag Foundation, and William K. Bowes Jr. Foundation (to T.J.N.). T.J.N. is a New York Stem Cell Foundation Robertson Neuroscience Investigator. E.E.E., A.C., and H.Y.Z. are investigators of the Howard Hughes Medical Institute.

## Author contributions

K.H., Y.S., E.E.E., and T.J.N. conceived the study and planned the experimental design. K.H., Y.S., D.K., A.N., and T.W. analyzed the data. K.H. and R.L. collected samples and performed experiments. R.M.B. and H.B.R. supported experiment planning and data analysis. K.H. and T.J.N. wrote the manuscript with input and feedback from all authors. H.Y.Z., A.C., E.E.E., and T.J.N. acquired funding for and supervised this work.

## Supplementary information

**Fig. S1:**
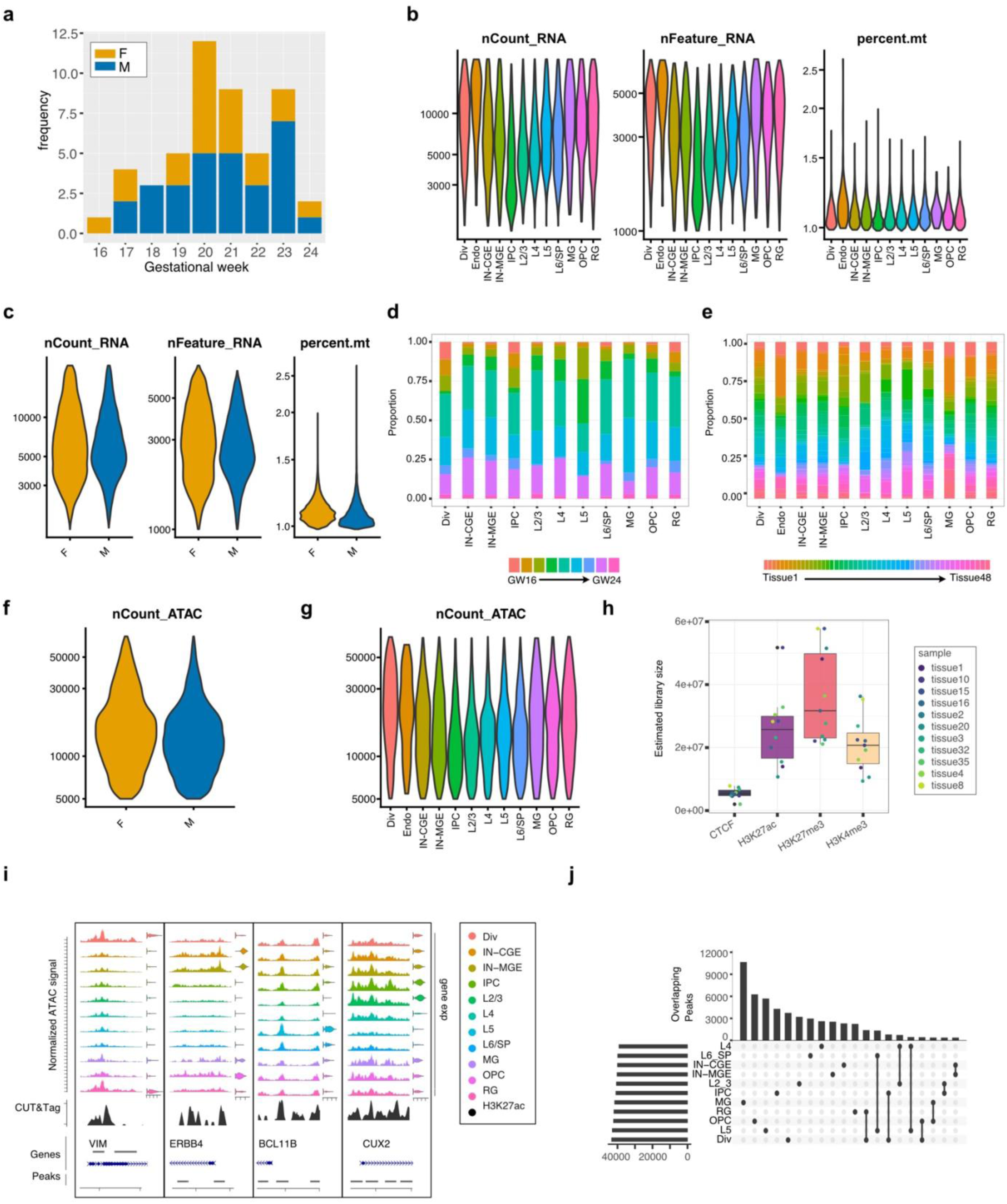
Quality control of single-nucleus Multiome and CUT&Tag samples. **a)** Histogram of samples by gestational week, by sex. **b)** Quality control (QC) by cluster of scRNA-seq data. **c)** QC by sex of scRNA-seq data, showing gene and UMI count, as well as mitochondrial gene percentage. **d)** Cluster proportion graph of sample age. GW = gestational week. **e)** Cluster proportion graph of donor. **f)** QC by sex of scATAC-seq data. **g)** QC by cell type of scATAC-seq data. **h)** Estimated CUT&Tag library size for all samples (n = 6 male, n = 5 female) of H3K27ac, H3K27me3, H3K4me3, and CTCF. **i)** Coverage plots of cluster markers VIM (radial glia progenitors), ERBB4 (inhibitory neurons), BCL11B (deep layer excitatory neurons), and CUX2 (upper layer excitatory neurons). Overlap of ATAC signal, H3K27ac signal, gene track, called ATAC peaks, and predicted peak-gene links are shown, with additional violin plots of gene expression by cell type. **j)** Upset plot of scE2G predicted CREs by cell type.

**Fig. S2:**
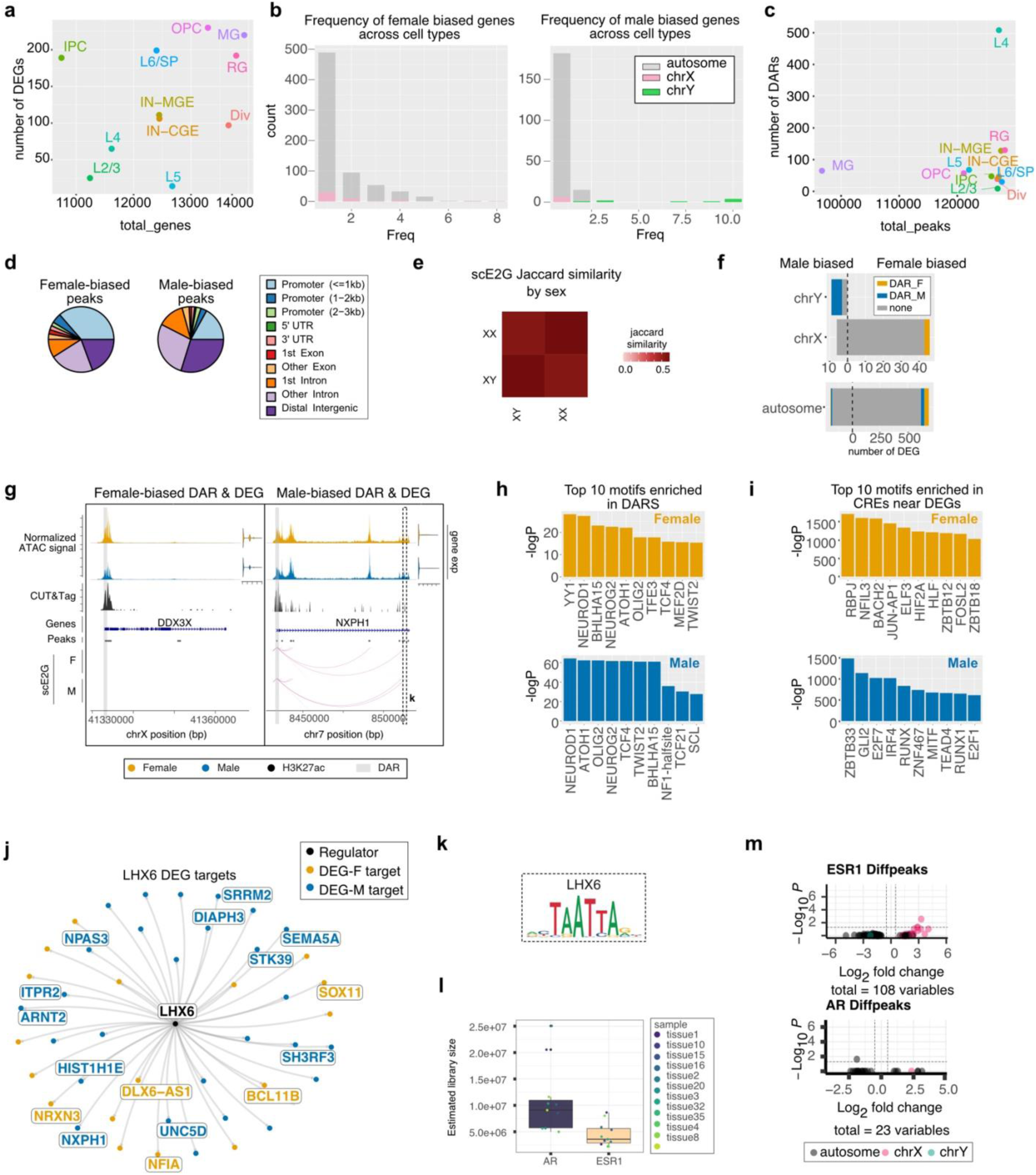
Characterizing sex biased gene expression, chromatin accessibility, motif enrichment, and gene regulatory networks. **a)** Scatter plot of total number of genes expressed (x-axis) by differentially expressed genes (y-axis) by cell type. **b)** Histogram of DEGs by number of cell types they are differentially expressed in. Grey is autosomal, pink is chrX, and green is chrY. **c)** Scatter plot of total number of accessible regions (x-axis) by DARs (y-axis) by cell type. **d)** Pie chart of annotation for female-biased and male-biased DARs. **e)** Correlation matrix of Jaccard similarity scores for CREs predicted for each sex by the scE2G pipeline. **f)** Bar plot of female-biased (right side) and male-biased (left side) genes. Colors indicate if the DEG has a DAR. **g)** Coverage plot of female biased gene *DDX3X* and male biased gene *NXPH1*, with normalized ATAC tracks and gene expression by sex, as well as H3K27ac CUT&Tag coverage and scE2G CRE prediction by sex. **h)** Top 10 motifs enriched in CREs near female biased (top, orange) or male biased (bottom, blue) DEGs. **i)** Top 10 motifs enriched in female biased (top, orange) or male biased (bottom, blue) DARs **j)** Regulon plot for LHX6 predicted targets that are also DEGs. Targets that are female-biased DEGs are orange (n = 18), targets that are male-biased DEGs are blue (n = 27). Total targets n = 467. **k)** LHX6 motif, seen at *NXPH1* promoter (g). **l)** Estimated library size for AR and ESR1 CUT&Tag across all samples. **m)** Sex-biased ESR1 and AR peaks. Peaks shown pass nominal significance threshold, where adjusted p-value is shown.

**Fig. S3:**
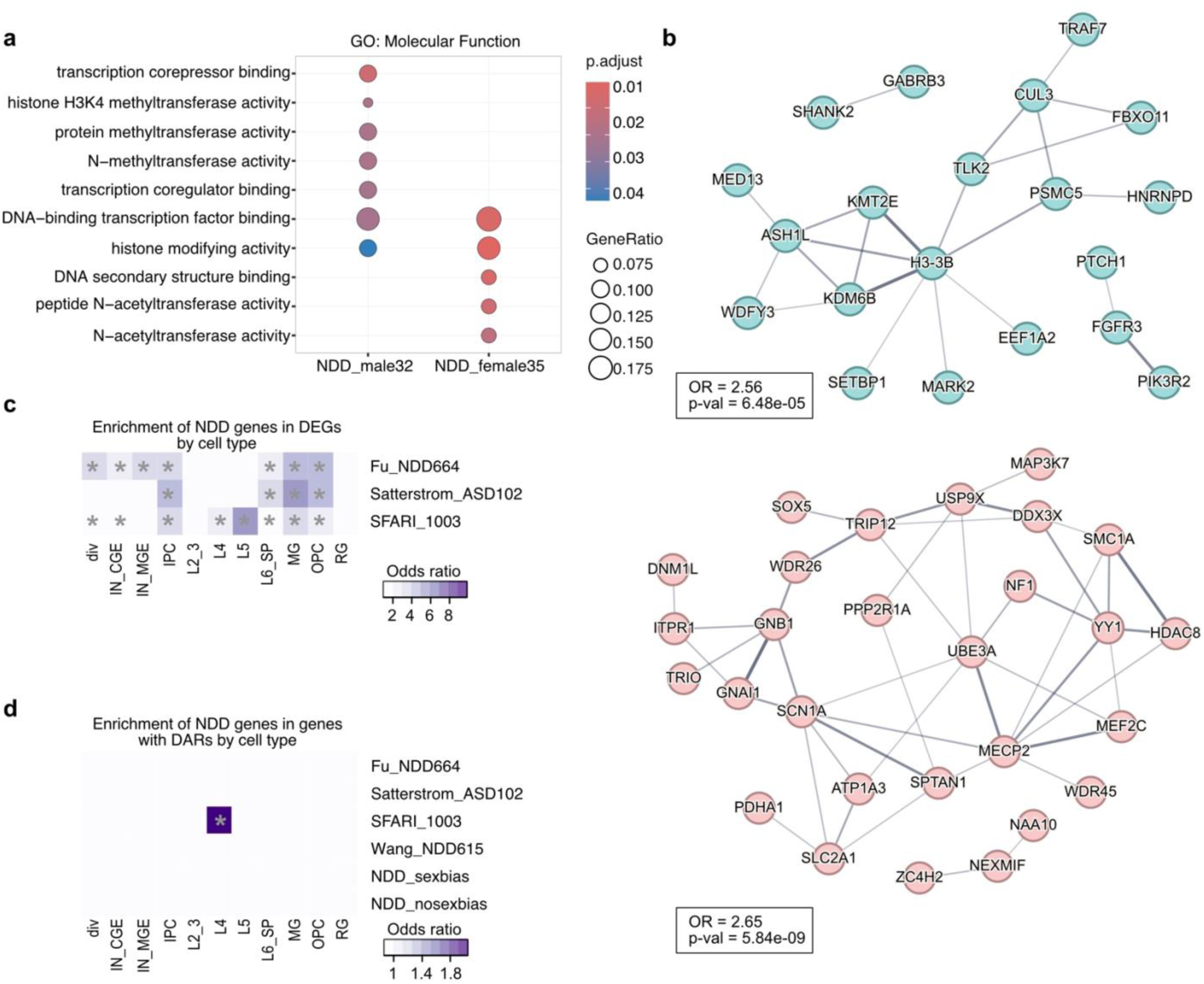
Characterizing sex-biased NDD genes. **a)** Dotplot of molecular function gene ontology for male biased NDD genes (SSD_male) and female biased NDD genes (SSD_female). Color represents BH corrected p-value. Dot size indicates gene ratio. **b)** STRING protein interaction network of male (left) and female (right) biased NDD genes. Line thickness indicates the strength of data support from the STRING v12.0 database. **c)** Enrichment of NDD genes from different studies in DEGs by cell type. * p < 0.01 (BH correction). **d)** Enrichment of NDD genes from different studies in genes with DARs by cell type. * p < 0.01 (BH correction).

## Supplementary table legends

**Supplementary table 1.** snMultiome sample metadata.

**Supplementary table 2.** Cluster markers for RNA assay.

**Supplementary table 3.** Map of consensus ATAC peaks from this study and Ziffra 2021.

**Supplementary table 4.** CTCF consensus CUT&Tag peaks (q < 0.01).

**Supplementary table 5.** H3K27ac consensus CUT&Tag peaks (q < 0.01).

**Supplementary table 6.** H3K4me3 consensus CU&Tag peaks (q < 0.01).

**Supplementary table 7.** H3K27me3 consensus CUT&Tag peaks (q < 0.01).

**Supplementary table 8-19.** scE2G predicted enhancer-gene interactions by cell type, using the multiome_powerlaw_v3 model.

**Supplementary table 20.** snRNA-seq differentially expressed genes between males and females by cell type (FDR < 0.05).

**Supplementary table 21.** snATAC-seq DARs between males and females by cell type (FDR < 0.05, log2FC >|1.25|). Positive log2FC = male-biased, negative log2FC = female-biased.

**Supplementary table 22.** scE2G predicted enhancer-gene interactions for bulk female cells, using the multiome_powerlaw_v3 model.

**Supplementary table 23.** scE2G predicted enhancer-gene interactions for bulk male cells, using the multiome_powerlaw_v3 model.

**Supplementary table 24.** Intersection of female-biased DARs (Supplementary table 20) with scE2G predicted enhancer-gene interactions in female cells (Supplementary table 21) to map predicted gene targets of DARs.

**Supplementary table 25.** Intersection of male-biased DARs (Supplementary table 20) with scE2G predicted enhancer-gene interactions in male cells (Supplementary table 22) to map predicted gene targets of DARs.

**Supplementary table 26.** HOMER Motif enrichment analysis in DARs (hypergeometric test, one-sided).

**Supplementary table 27.** HOMER Motif enrichment analysis in CREs near DEGs (hypergeometric test, one-sided).

**Supplementary table 28.** scMEGA gene regulatory network analysis.

**Supplementary table 29.** ESR1 consensus CUT&Tag peaks (q < 0.01), with annotation of nearest neighbor gene.

**Supplementary table 30.** AR consensus CUT&Tag peaks (q < 0.01), with annotation of nearest neighbor gene.

**Supplementary table 31.** Sex-biased NDD genes NDD_male32 and NDD_female35, and genes with no sex bias NDD_both97 (two-sided Fisher exact test, BH correction). Genes were determined for integrated NDD as well as separate ASD and DD cohorts. The sex-biased genes from the integrated NDD analysis were used for further comparison in this study.

